# A two-step metagenomics approach for prey identification from the blood meals of common vampire bats (*Desmodus rotundus*)

**DOI:** 10.1101/2021.12.06.470776

**Authors:** Physilia Ying Shi Chua, Christian Carøe, Alex Crampton-Platt, Claudia Sarai Reyes-Avila, Gareth Jones, Daniel G. Streicker, Kristine Bohmann

## Abstract

The feeding behaviour of the sanguivorous common vampire bat (*Desmodus rotundus*) facilitates the transmission of pathogens that can impact both human and animal health. To formulate effective strategies in controlling the spread of diseases, there is a need to obtain information on which animals they feed on. One DNA-based approach, shotgun sequencing, can be used to obtain such information. Even though it is costly, shotgun sequencing can be used to simultaneously retrieve prey and vampire bat mitochondrial DNA for population studies within one round of sequencing. However, due to the challenges of analysing shotgun sequenced metagenomic data such as false negatives/positives and typically low proportion of reads mapped to diet items, shotgun sequencing has not been used for the identification of prey from common vampire bat blood meals. To overcome these challenges and generate longer mitochondrial contigs which could be useful for prey population studies, we shotgun sequenced common vampire bat blood meal samples (n=8) and utilised a two-step metagenomic approach based on combining existing bioinformatic workflows (alignment and *de novo* mtDNA assembly) to identify prey. Further, we validated our results to detections made through metabarcoding. We accurately identified the common vampire bats’ prey in seven out of eight samples without any false positives. We also generated prey mitochondrial contig lengths between 138bp to 3231bp (mean=985bp, SD=981bp). As we develop more computationally efficient bioinformatics pipelines and reduce sequencing costs, we can expect an uptake in metagenomics dietary studies in the near future.

## INTRODUCTION

The common vampire bat (*Desmodus rotundus*) is one of three extant species of vampire bats (Chiroptera; Phyllostomidae; Desmodontinae) native to Latin America (Greenhall et al., 1983). It has an obligatory sanguivorous diet and feeds on vertebrate blood by biting its prey. The common vampire bat is therefore primed to facilitate cross-species transmission of pathogens such as *Bartonella* (Bai et al., 2012), hemoplasmas (Volokhov et al., 2017), and trypanosomes (Hoare, 1965). Common vampire bats are also the primary reservoir of the rabies virus in much of Latin America (Schneider et al., 2009). Rabies is a lethal zoonotic disease, killing thousands of livestock annually and causing sporadic outbreaks in human populations where bats routinely feed on humans (Schneider et al., 2009). Land-use change from forest to livestock pastures has provided the common vampire bats with an abundant and accessible source of mammalian prey, leading to population growth and range expansion (Delpietro et al., 1992; Lee et al., 2012; Streicker & Allgeier, 2016). These bats are also generalist feeders, able to feed not just on livestock but also on wildlife including marine species such as sea lions and penguins (Luna-Jorquera and Culik 1995; Catenazzi and Donnelly 2008). As their distribution continues to respond to climate change, feeding patterns of the common vampire bats can be expected to change accordingly (Hayes and Piaggio 2018). Hence, knowing the diet of the common vampire bat and thereby, their potential routes of disease transmission, is a necessary step to control the spread of pathogens in a cost-effective and efficient manner. One way of determining what taxa they feed from and can potentially transmit pathogens to, is by identifying prey species in vampire bat blood meals (Bohmann et al., 2018; Greenhall, 1988).

The different types of methods used for the identification of common vampire bat prey include field observations (Catenazzi & Donnelly, 2008; Greenhall, 1988), camera traps (Calfayan et al., 2018; Galetti et al., 2016; Zortéa et al., 2018), precipitation tests to visualise antibody-antigen complexes (Greenhall, 1970), and stable isotope analysis (Catenazzi & Donnelly, 2008; Streicker & Allgeier, 2016; Voigt & Kelm, 2006). Field observations are challenging as bats are nocturnal (Tournayre et al., 2021), precipitation tests are labour intensive, and stable isotope analysis does not give species resolution (reviewed in (Carter et al., 2021)). DNA-based methods such as metabarcoding are faster and more precise, allowing for species identification of prey and simultaneous retrieval of common vampire bat population structure (Bohmann et al., 2018). However, amplification of vertebrate prey from blood meals can be challenging due to PCR inhibitors present in blood (Akane et al., 1994), and the co-amplification of vampire bat DNA which could prevent the detection of fragmented prey DNA present in lower copy numbers (Bohmann et al., 2018). To overcome these challenges, blocking primers could be used to reduce the amplification of predator DNA (Deagle et al., 2009; Vestheim & Jarman, 2008). However, the design of predator-blocking primers can be made difficult by the lack of DNA reference sequences for prey species, and for the common vampire bat in particular by high intraspecific variation in the common vampire bat mitochondrial genome (Bohmann et al., 2018).

Another DNA-based method that could be used to identify the prey species in the common vampire bat blood meal diet is metagenomics. Metagenomics is where the DNA extracted from samples are shotgun-sequenced without target enrichment of specific markers (Noonan et al., 2005). This has the caveat that sequencing costs are at least ten times more expensive than metabarcoding (Chua et al. 2021). This currently limits the number of samples that can be sequenced using this approach. However, shotgun sequencing overcomes the need to select specific vertebrate primers and can simultaneously retrieve prey and predator mitochondrial DNA, predator gut microbiome and gut parasites (Ang et al., 2020; Bon et al., 2012; Paula et al., 2015, 2016; Srivathsan et al., 2015, 2016). This maximises the amount of information that can be retrieved within one round of sequencing, without the need to carry out additional lab work as is required for metabarcoding using multiple primers.

Despite the advantages of shotgun sequencing, bioinformatics analyses of these metagenomics data can be challenging where false negatives and positives are often an issue (Chua et al. 2021; Paula et al. 2016). Two main strategies can be used to identify metagenomic reads, namely alignment-based and assembly-based approaches. The alignment-based approach is where reads are mapped to a DNA reference database for identification (Zhang et al., 2000), and the resulting mapped reads are identified to the Last Common Ancestor (LCA) for identification (Huson & Weber, 2013). However, such identifications are dependent on the completeness of the reference database used (Chua et al., 2021; Gómez□Rodríguez et al., 2015). In diet studies, a low proportion of reads (between 0.0001% to 0.009%) are typically mapped to diet items when using this approach (Alberdi et al., 2018; Chua et al., 2021; Srivathsan et al., 2015, 2016). The reliance on a low proportion of reads to inform results increases the risks of false positives (Paula et al. 2016).

To increase the proportion of informative reads mapped to diet items and generate longer reads which could be useful for further analysis on prey population structure, a *de novo* assembly-based approach of assembling mitochondrial DNA (mtDNA) contigs can be carried out using dedicated assemblers (Dierckxsens et al., 2017). However, this is traditionally only used for mtDNA assembly of a single known organism as it requires the selection of an appropriate input reference seed file used for assembly. These reference seed files are usually barcode markers of the known organisms which are used for extending the reads to generate longer mtDNA contigs. In known-mixed template samples, assemblies are limited to the taxa with the highest proportion of reads in the sample (reviewed in (Sharpton, 2014)). The assembly of low abundance sequences will be fragmented if the sequencing depth is too low. This can be problematic in diet studies given that predator sequences would overwhelm the proportion of prey sequences. Conserved regions shared between prey and predator could also lead to the assembly of chimeric sequences (Bon et al., 2012).

In animal dietary studies, shotgun sequencing has been used to identify plants in the diets of herbivores (Chua et al., 2021; Srivathsan et al., 2015, 2016), arthropod prey from arthropod predators (Paula et al., 2015, 2016), and vertebrate prey from vertebrate predators (Bon et al., 2012). In common vampire bat studies, shotgun sequencing of faeces, saliva, or rectal swabs has been used to identify the bat’s genomic adaptations to sanguivory (Mendoza et al., 2018), profile its gut microbiome (Mendoza et al., 2018), characterise its viral communities (Bergner et al., 2019; Bergner, Orton, Benavides, et al., 2020), and to assemble the genomes of common vampire bat viruses (Bergner, Orton, & Streicker, 2020). However, identification of prey from common vampire bat blood meal samples has not been carried out using the metagenomics shotgun sequencing approach.

Here, we use shotgun sequencing for the first time to identify prey from common vampire bat blood meal samples. To overcome the challenges associated with each of the shotgun sequencing approaches outlined above, we demonstrate the advantages of combining both the alignment and assembly-based approaches in a stepwise manner to reduce the limitations associated with each approach. To validate our results, we verified our metagenomic prey identification with the metabarcoding results obtained from the same samples by Bohmann et al. (2018).

## MATERIALS AND METHODS

### Sample information

The eight blood meal samples from common vampire bats (*Desmodus rotundus*) used were collected in Peru between 2009 and 2013 at four sites across two ecoregions; Amazon (MDD130) and Pacific coast (LMA4, LMA6, and LMA10) (Supplementary Table S1). Four samples were collected each from the Amazon (sample 25, 54, 116, and 121) and the coastal ecoregions (sample 29, 70, 90, and 94). The blood meal collection followed the procedures outlined in (Streicker & Allgeier, 2016). From the metabarcoding results outlined in Bohmann et al. (2018), the prey taxa of the eight common vampire bat blood meal samples were identified as chicken (*Gallus gallus*), cow (*Bos taurus*), donkey (*Equus asinus*), horse (*Equus caballus*), pig (*Sus scrofa*), and the South American tapir (*Tapirus terrestris*) (Fig. 1).

**Figure 1:**
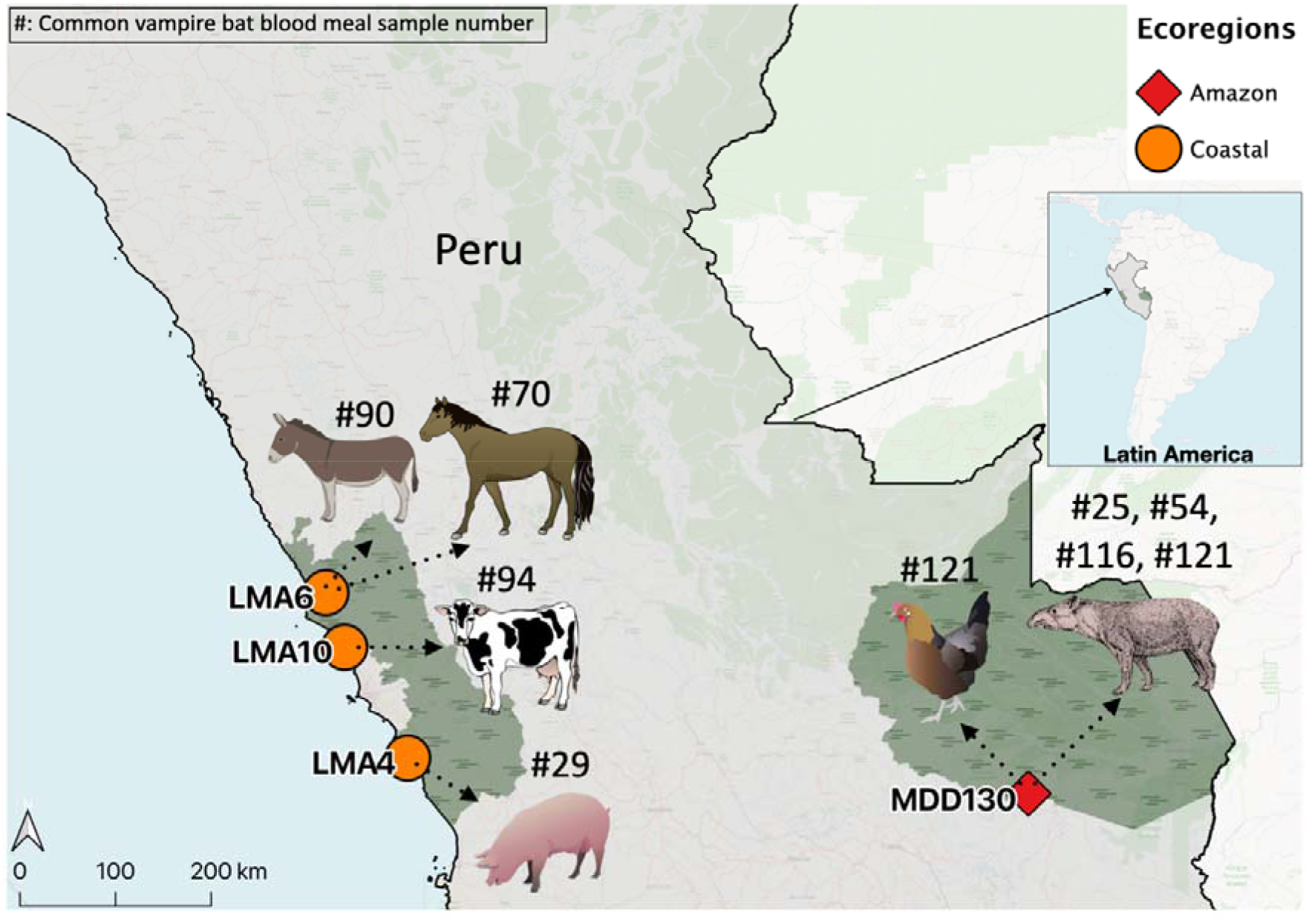
Collection sites of eight common vampire bat (*Desmodus rotundus*) blood meal samples from two ecoregions in Peru, showing the identity of the prey taxa derived from metabarcoding analysis (Bohmann et al., 2018). Map created in *QGIS version 3*.*12*. Some of the elements included in the figures were obtained and modified from the Integration and Application Network, University of Maryland – Center for Environmental Science (https://ian.umces.edu/symbols/), and BioRender.com. Image of tapir from Foresman (2007).

### Metagenomics laboratory workflow

We used the extracted DNA from the eight blood meal samples that were previously extracted as described in Bohmann et al. (2018). DNA extracts were fragmented using a Diagenode Bioruptor (Diagenode) using a program of eight cycles with 15 seconds on and 90 seconds off targeting a fragment size of 500 bp. 32 μL of fragmented DNA was used to generate Illumina shotgun sequencing libraries using the blunt-end single-tube library preparation protocol (Carøe et al., 2018) with modifications from (Mak et al., 2017). The libraries were purified using SPRI bead purification according to (Rohland & Reich, 2012). Specifically, 100 μL of bead solution was added to each library (60 μL), incubated for 5 minutes, washed twice in 80% ethanol and eluted in 30 μL of 10 mM Tris-HCl by heating to 40°C for 5 minutes. Libraries were evaluated with quantitative PCR (qPCR), 480 Lightcycler 2x qPCR mastermix (Roche) in 10 μL reactions, with 0.2μM primer IS7 and IS8 (Meyer & Kircher, 2010), and 1 μL of 10x diluted library. Based on cyclic threshold values, libraries were given 7 to 11 PCR cycles for index PCR, using full-length Illumina primers with indexed P7 adapters. This was done using 10 μL library in a 50 μL PCR reaction consisting of 0.25 mM dNTP (Invitrogen), 0.2 mM forward and reverse primer, 0.1 U/mL Taq Gold polymerase (Applied Biosystems), 1x Taq Gold buffer (Applied Biosystems), 2.5 mM MgCl_2_ (Applied Biosystems), and 0.8 mg/mL BSA (New England Biolabs). PCR consisted of 10 minutes denaturation and activation at 95°C, followed by 7 to 11 cycles of 30 seconds at 95°C, 30 seconds at 60°C, and 1 minute at 72°C, followed by a final extension at 72 °C for 5 minutes before cooling to 4°C. Libraries were purified using MinElute spin columns (Qiagen). Purified libraries were quantified on an Agilent 2100 bioanalyzer (Agilent technologies) before equimolar pooling. Sequencing was carried out on one lane of an Illumina 2500 Hiseq instrument (Illumina Inc.) using 125 cycle chemistry in paired-end (PE) mode at the GeoGenetics Sequencing core, University of Copenhagen, Denmark.

### Metagenomics bioinformatics workflow

Between ∼17 and ∼36 million paired-end (PE) reads were generated per blood meal sample. Adapter removal, quality trimming of sequences with Phred quality score less than 30, and removal of reads shorter than 85 base pairs (bp) were carried out with *Trim galore v0*.*5*.*0* (Andrews et al., 2015). This cut-off length at two-thirds of the sequenced reads was introduced to reduce the rates of false-positive identification in downstream analysis, while keeping most true-positive reads (Chua et al., 2021; Srivathsan et al., 2015, 2016). *FastQC v0*.*11*.*9* was used for quality checks before and after filtering (Andrews, 2010). For each sequenced blood meal sample, we used the *Burrows-Wheeler Alignment* software with the *Maximal Exact Matches* algorithm *v0*.*7*.*17 (BWA-MEM)* and *Sequence Alignment/Map v1*.*9 (SAMtools)* software to map and align PE reads to the common vampire bat genome downloaded from NCBI GenBank (RefSeq assembly accession: GCF_002940915.1) (downloaded 27.04.20). Aligned reads mapped to the common vampire bat genome were subsequently removed from each of the sequenced blood meal samples (Li, 2013; Li et al., 2009). Using the *Browser Extensible Data v2*.*29 (BEDtools)* software (Quinlan & Hall, 2010), we converted the BAM files generated in the common vampire bat sequence-removal step to fastq files, using the *bamToFastq* function, for downstream analysis.

#### Prey identification

For the identification of common vampire bat blood meal prey taxa, we used a two-step approach. In the first step, we used an alignment method, in which we carried out the Basic *Local Alignment Search Tool (BLAST*) to map reads to reference data. For the second step, based on the *BLAST* results, we used a seed (COI barcode or whole mtDNA) corresponding to the identity of the prey to carry out *de novo* assembly of prey mtDNA contigs. The *de novo* mtDNA contig assembly step was also used to retrieve any additional identification of prey from blood meal samples that had no results from the *BLAST-*alignment step.

##### Step 1: *BLAST*-alignment step

We generated a reference database by downloading taxonomically informative barcodes consisting of metazoan mitochondrial cytochrome oxidase subunit 1 (COI) sequences from the Barcode of Life Data System v4 (*BOLD*) (473,748 sequences forming 38,618 BINS, representing 33,299 metazoan species downloaded 27.04.20) (Ratnasingham & Hebert, 2007). *MEGABLAST* searches for the sequenced bloodmeal PE reads were conducted against the generated COI barcode reference database (word size = 28, percentage identity = 98%) (Camacho et al., 2009; Srivathsan et al., 2015, 2016). We used the *bold* package in R with a custom R script *BOLD_taxID* to retrieve taxonomy classification details for each BOLD BIN in the generated COI barcode reference database (retrieved 29.04.20). For taxonomic assignment of the common vampire bat blood meal sequences to determine prey species, we used a custom R script *BOLD_readsidentifier* with the following filtering parameters of 98% sequence identity and 85 bp overlap of a given read with the COI barcode (Srivathsan et al., 2015, 2016). Only sequences identified as belonging to the classes Aves or Mammalia were kept. Following the Lowest Common Ancestor algorithm (Huson & Weber, 2013), we obtained species-level identification for a given read if the *BLAST* hit was to one species, genus-level if it matches to two or more species from one genus, and family level if it matches to two or more genera from the same family. We only retained identifications at a given taxonomic hierarchy where there were no conflicts in identification between the forward and reverse reads (Srivathsan et al., 2015, 2016). For samples with more than one species identified, we only used the identity of the species with the highest number of reads matched. This is supported by the expectation that vampire bats feed on a single individual per night, thus secondary reads are likely to represent false positives (Greenhall, 1988.)

##### Step 2: *De novo* mtDNA contig assembly step

For the *de novo* mtDNA contig assembly step, we used the organelle assembler and heteroplasmy caller software *NOVOPlasty v3*.*8*.*3* (Dierckxsens et al., 2016) to attempt assemblies of prey mtDNA contigs. *NOVOPlasty* is a seed-based assembler, where assembly is initiated by a reference seed file. Reference seed files of specific prey species identified from each blood meal sample from the initial *BLAST*-alignment step were used as inputs for mtDNA contig assembly. If more than one prey species was identified per blood meal sample from the *BLAST*-alignment step, we used reference seed files of the species with the highest number of reads mapped to the COI barcode. For blood meal samples with no prey identified from *BLAST*, we used reference seed files of all prey species found in the four sites that have been identified as prey using metabarcoding (Bohmann et al., 2018). Reference seed files used were either species-specific COI barcodes retrieved from BOLD (downloaded 17.06.20) or whole mtDNA retrieved from NCBI GenBank (downloaded 27.08.20) (Supplementary Table S2 – S4). We carried out assembly with a K-mer size of 39 starting with a species-specific COI barcode as seed, and decreasing the K-mer size to 27 if no contigs were assembled. We changed the input seed files to species-specific mtDNA if no contigs were assembled after decreasing K-mer size to 27. Assembled mtDNA contigs were checked by using the *web BLAST blastn suite*, to obtain the closest match (Madden, 2013). Contigs were also manually checked for alignment to reference seed using *Geneious Prime v2020*.*2* (https://www.geneious.com). To ensure the accuracy of the *de novo* assembly step, we attempted mtDNA contig assembly with seed files of species not identified as prey for all blood meal samples with K-mer size 39 (COI barcode of all species in Supplementary Table S2, and mtDNA of *Tapirus terrestris*). Any contig(s) assembled were checked by uploading the contig sequences to the *web BLAST blastn* suite for sequence identity (Supplementary Table S5). Only the closest matched species in terms of query cover and percent identity (> 90%) were kept.

After these two steps, the metagenomics outputs from this study were compared with the metabarcoding results from Bohmann et al. (2018) to check for any discrepancies between the two approaches.

## RESULTS

Reads mapped to the common vampire bat genome (*Desmodus rotundus)* made up 48.1% to 98.7% of PE reads generated per blood meal sample (∼16 million to ∼60 million reads, average 48 million reads, SD = 13 million reads). After removal of common vampire bat sequences, 1.3% to 51.9% of reads remained which includes sequences from the prey, the common vampire bat’s gut microbiome, and gut parasites (∼67,000 to ∼17 million reads per sample, average 6.7 million reads, SD = 5.9 million reads) (Table 1).

**Table 1:** Proportion of metagenomic reads belonging to the common vampire bat (*Desmodus rotundus*) and proportion of reads assembled (common vampire bat-removed sequences) from the *de novo* mtDNA contig assembly step for prey identification.

| Samples | Total reads | Bat sequences | Proportion of bat sequences | Bat-removed reads for downstream analysis | Assembled mtDNA reads | Proportion of reads assembled for prey identification |
| --- | --- | --- | --- | --- | --- | --- |
| 25 | 57782738 | 55744368 | 96.5% | 2038370 | 40 | 0.00196% |
| 29 | 61585369 | 60726115 | 98.6% | 859254 | 50 | 0.00582% |
| 54 | 30704614 | 21223068 | 69.1% | 9481546 | 3478 | 0.03668% |
| 70 | 61417998 | 51802288 | 84.3% | 9615710 | 48 | 0.00050% |
| 90 | 56874109 | 55157869 | 97.0% | 1716240 | 3500 | 0.20393% |
| 94 | 31315790 | 19574502 | 62.5% | 11741288 | 3010 | 0.02564% |
| 116 | 50267627 | 49596878 | 98.7% | 670749 | 0 | 0.00000% |
| 121 | 33684838 | 16217550 | 48.1% | 17467288 | 10 | 0.00006% |

### Prey identification

In the *BLAST*-alignment step, *BLAST* searches against the COI database yielded between 1 and 36 reads mapped to Mammalia or Aves (<0.0001% - 0.003% of vampire-bat removed reads, <0.0001% - 0.0001% of total sequenced reads) for five of the blood meal samples (samples 54, 70, 90, 94 & 121). Three blood meal samples did not have any reads mapped to the COI barcode for either Mammalia or Aves (samples 25, 29, & 116) (Supplementary Table S6). These three bloodmeal samples each contained more than 95% of common vampire bat sequences prior to common vampire bat sequence removal. For the five blood meal samples with reads mapped to the COI barcode, the majority of the reads were assigned species-level identification (57.6%), 37.3% of the reads were assigned to genus, 3.4% of the reads assigned to family, and 1.7% of the reads assigned to order. For these five blood meal samples, four had reads assigned to only one species except for sample 94 with five species identified (1 read: *Bison bonasus*, 1 read: *Bos grunniens*, 2 reads: *Bos indicus*, 1 read: *Bos primigenius*, 15 reads: *Bos taurus*). We only kept the identity of the species with the highest number of reads matched, which was *Bos taurus*. The species identity of the prey found in the five blood meal samples were; *Tapirus terrestris* (sample 54), *Equus caballus* (sample 70), *Equus asinus* (sample 90), *Bos taurus* (sample 94), and *Gallus gallus* (sample 121).

For *de novo* assembly of mtDNA contigs, the proportion of reads assembled was between 0.0006% - 0.2% of vampire-bat removed reads (Table 1). The average length of mtDNA contigs assembled was 985bp (SD=981bp), with the smallest being 138bp from sample 121 and the largest 3231bp from sample 94 (Table 2). Most samples had only one contig assembled except for samples 54 and 94, with two and three contigs assembled respectively. The species identity of the prey from the assembled mtDNA contigs corresponded to the *BLAST*-alignment results for all five blood meal samples (samples 54, 70, 90, 94 & 121). We also retrieved prey information for two additional samples (samples 25 & 29) from the *de novo* mtDNA contig assembly which had no identification after the first *BLAST-*alignment step. The identity of the prey from these two blood meal samples were *Tapirus terrestris* for sample 25, and *Sus scrofa* for sample 29. All identified prey were identified to the species level. We did not retrieve prey identification for sample 116 after the *BLAST*-alignment and *de novo* mtDNA contig assembly steps (Fig. 2).

**Table 2:** Prey identity of *de novo* assembled mtDNA contigs for each common vampire bat (*Desmodus rotundus*) blood meal shotgun-sequenced sample with query cover and percent identity score.

| Sample | Contig length (Bp) | Closest Match | Query Cover | Percent identity |
| --- | --- | --- | --- | --- |
| 25 | 684 | *Tapirus terrestris* | 100% | 91.23% |
| 29 | 166 | *Sus scrofa* | 92% | 98.70% |
| 54 | 1152 | *Tapirus terrestris* | 100% | 90.89% |
|  | 770 | *Tapirus terrestris* | 97% | 91.71% |
| 70 | 268 | *Equus caballus* | 98% | 94.70% |
| 90 | 256 | *Equus asinus* | 100% | 94.53% |
| 94 | 3231 | *Bos taurus* | 100% | 94.15% |
|  | 2380 | *Bos taurus* | 100% | 94.33% |
|  | 802 | *Bos taurus* | 100% | 94.39% |
| 121 | 138 | *Gallus gallus* | 100% | 100% |

**Figure 2:**
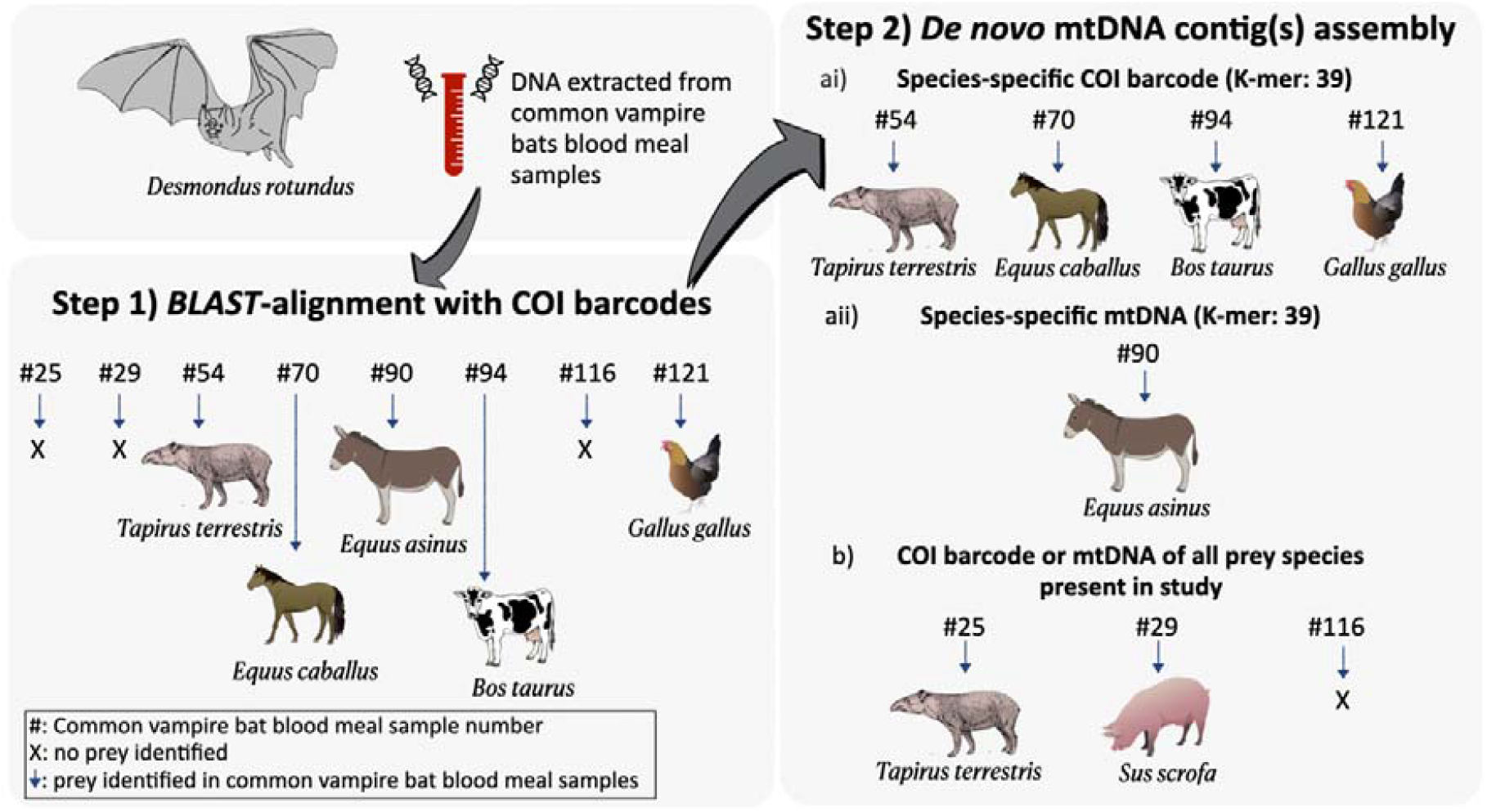
Prey identification of common vampire bat (*Desmondus rotundus*) blood meal shotgun-sequenced samples using a two-step metagenomics approach. In the first *BLAST*-alignment step, metagenomic reads were mapped to COI barcodes using *BLAST*. Second, *de novo* mtDNA assembly of prey contigs was carried out with Novoplasty using ai) seeds from COI barcode or aii) mitochondrial DNA (mtDNA) of prey identified from the *BLAST*-alignment step, and b) seeds from COI barcode or mtDNA of all prey species identified from the metabarcoding analysis presented in Bohmann et al. (2018). Some of the elements included in the figures were obtained and modified from the Integration and Application Network, University of Maryland – Center for Environmental Science (https://ian.umces.edu/symbols/), and BioRender.com. Image of tapir from Foresman (2007). Common vampire bat image credit: Megan Griffiths.

When comparing the metagenomics outputs with the metabarcoding analysis presented in Bohmann et al. (2018), there was a 100% congruence in the identity of the prey detected using metagenomics for all samples, with no false positives. However, metagenomics failed to identify *Tapirus terrestris* found in the blood meals of samples 116 and 121, leading to a 22% false negative detection rate (Fig. 3).

**Figure 3:**
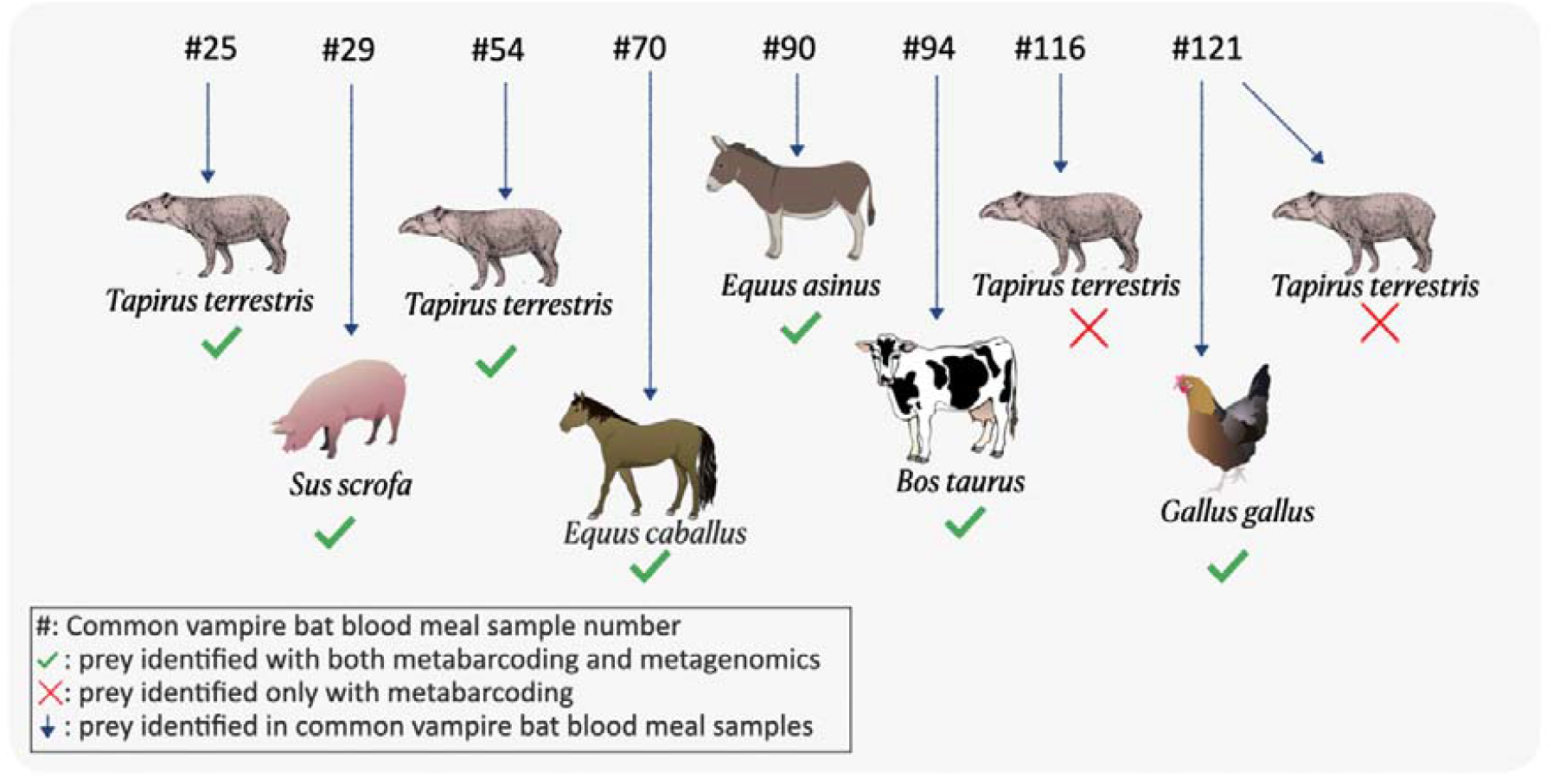
Prey identification of common vampire bat blood meal samples using shotgun metagenomics as compared to metabarcoding (Bohmann et al., 2018). Green ticked symbols signify consensus between both high-throughput sequencing (HTS) approaches. Red crossed symbols signify prey taxa identified using metabarcoding but not identified using metagenomics. Some of the elements included in the figures were obtained and modified from the Integration and Application Network, University of Maryland – Center for Environmental Science (https://ian.umces.edu/symbols/), and BioRender.com. Image of tapir from Foresman (2007).

## DISCUSSION

In this study, we demonstrated how metagenomics can be used to identify common vampire bat prey in blood meals. We utilised a two-step strategy, combining both the alignment and assembly approaches to obtain prey identification. These metagenomic prey identifications were subsequently compared to metabarcoding results for validation (Bohmann et al., 2018).

In the *BLAST*-alignment approach, the proportion of reads mapped to prey is similar to other metagenomics studies (Alberdi et al., 2018; Chua et al., 2021; Srivathsan et al., 2015, 2016). Despite the low proportion of reads mapped, we achieved good species resolution at 57.6%, which is higher than these recent metagenomic diet studies. However, in samples with a high proportion of vampire bat DNA (> 95% in samples #25, #29, and #116), we were unable to map any reads to prey. This could be due to the extremely low copy numbers and fragmented nature of prey DNA present. The completeness of the reference database used for the matching of short reads could also have resulted in these missing identifications (Chua et al., 2021). Without carrying out the second step *de novo* mtDNA contig assembly approach, stopping at this step of the bioinformatics analysis would result in missing prey identifications for two samples (#25 and #29).

From the *de novo* mtDNA contig assembly approach, a higher proportion of reads were assembled as compared to the *BLAST*-alignment step (0.0006% - 0.2%). This is expected because reads were only mapped to the COI barcode in the *BLAST*-alignment step while in the *de novo* mtDNA contig assembly step, longer mtDNA contigs were generated. The resolution of prey identified using this approach was at 100% species-level, which is another huge advantage of combining the two approaches. Even though the application of the *de novo* mtDNA contig assembly step for eukaryote mtDNA assembly can be problematic due to complex genomes and low abundance (Azam & Malfatti, 2007), we managed to retrieve prey identification for most samples except for one (#116), which had the highest proportion of common vampire bat sequences (98.7%). The high abundance of common vampire bat sequences found in blood meals could have drowned out the DNA sequences belonging to the prey during sequencing. This feeds into the limitation of the *de novo* mtDNA contig assembly step, where assemblies are limited to taxa with a high proportion of reads in a sample (Sharpton, 2014). This could also be the reason why we were only able to detect only one prey, *Gallus gallus* and not *Tapirus terrestris*, in the blood meal of sample 121.

Another concern was the selection of appropriate reference seed files for *de novo* mtDNA contig assembly. The selection of inappropriate reference seed could lead to Type I errors resulting in false positives and inaccurate determination of diet. However, when we tested for the accuracy of *de novo* mtDNA assembly by using reference seed files from other species, eg: *Equus asinus* for *Tapirus terrestris* in sample 54, the assembler chosen in our study managed to accurately assemble mtDNA contigs belonging to the identified prey. Based on our samples and the assembler used, the selection of input reference seed file did not result in any false positives but further tests should be carried out on larger sample sizes with a variety of reference seed files. When using other types of assemblers requiring input reference seed files, the choice of reference seed file utilised should be informed by first carrying out the *BLAST*-alignment approach and validated with metabarcoding data to prevent inaccurate identifications.

When we compared our results to previous metabarcoding analyses carried out on the same samples (Bohmann et al., 2018), we were able to accurately identify common vampire bat blood meal prey from seven out of the eight samples. For sample 121 in which the bat fed on two prey (*Gallus gallus* and *Tapirus terrestris*), we only managed to identify *Gallus gallus*. The drawback of metagenomics in common vampire bat diet analyses is that not all prey can be identified and sequencing costs are at least 10 times more than for metabarcoding (Chua et al., 2021). Based on sequencing costs calculated for this study, shotgun sequencing eight samples costs €2730 as compared to only €135 for metabarcoding 118 samples. Given that metabarcoding is cheaper, it will remain the go-to technique when it comes to molecular diet profiling of animals. The current costs of shotgun sequencing mean that published metagenomic animal dietary studies are typically limited to only a few individuals (Alberdi et al., 2018; Bon et al., 2012; Chua et al., 2021; Paula et al., 2016; Srivathsan et al., 2015, 2016). As such, bioinformatics procedures are still in their infancy for analysing metagenomics dietary datasets and pipelines used differ based on the type of diet, and the taxonomic class of the predator in comparison to its diet. Our strategy based on combining two existing bioinformatic workflows; the alignment and assembly-based approaches, can help to advance metagenomic dietary research and open doors for future study. The strategy outlined here is particularly useful in scenarios where information is absent about the diet or the genetic make-up of the collected samples, with the caveat that this two-step approach could be more computationally intensive and additional streamlining is needed to optimise performance. Nevertheless, this two-step metagenomics approach completely removes false-positives and minimises false-negatives, which is an important challenge to overcome when working with metagenomic datasets.

## FUTURE OUTLOOK

To fully utilise the potential of metagenomics, improvements in bioinformatic procedures are still required to optimise data analyses and to make the most of all sequenced data such as i) increasing the proportion of informative reads for diet analyses, ii) streamlining bioinformatics pipelines to reduce computational requirements, and iii) analysing the large amount of generated data to prevent “data-wastage”. For example in our study, a large proportion of metagenomic reads are mapped to the predator itself (48.1% to 98.7%) which could be used for population genetic studies of the common vampire bat. Other types of information such as gut microbiome composition and parasites could also be analysed from the same metagenomic data (Srivathsan et al. 2016; Srivathsan et al. 2015; Mendoza et al. 2018; Bergner et al. 2021), potentially giving a more holistic view of the common vampire bat ecology without the need to carry out additional sequencing as with metabarcoding. Additionally, existing metagenomic data that has not been analysed for diet can also be repurposed and analysed for diet studies without the need to carry out additional sequencing. As we continually make progress in reducing sequencing costs and developing more computationally efficient bioinformatics pipelines, we can expect there to be a shift towards more metagenomics dietary studies in the future.

## AUTHOR CONTRIBUTIONS

PYSC: Conceived the study, carried out bioinformatics analysis, data analysis and visualisation, and wrote the manuscript;

CC: Carried out laboratory analysis and contributed to the writing of the manuscript;

AP: Wrote the custom R scripts used in the BLAST-alignment approach and supervised data analysis;

CSRA: carried out the removal of vampire-bat sequences and contributed to the writing of the manuscript;

GJ: Contributed to the writing of the manuscript;

DS: Collected samples and contributed to the writing of the manuscript;

KB: Conceived and supervised the study, carried out laboratory analysis, and contributed to the writing of the manuscript

## ACKNOWLEDGMENTS

We thank the staff at GeoGenetics Sequencing core, University of Copenhagen, Denmark, for sequencing. PYSC was supported by the European Union Horizon 2020 research and innovation programme under grant agreement No 765000, H2020 MSCA-ITN-ETN Plant.ID network. CC was funded by the Carlsberg Foundation (CF18-1110;”Archives”). Field work was supported by the US National Science Foundation (DEB-1020966). DS was funded by a Wellcome Senior Research Fellowship (217221/Z/ 19/Z).

## DATA ACCESSIBILITY

Sequencing data, assembled contigs, BLAST outputs, and scripts used for data generation is available from the Dryad Digital Repository, XXXXX, upon acceptance of the manuscript. It is currently available at the University of Copenhagen Electronic Research Data Archive (UCPH ERDA) Digital Repository, https://erda.ku.dk/archives/1d5f6654c1bbfd5a0485d91ec8fa16c1/published-archive.html

