## Supplementary Table for "A two-step metagenomics approach for prey identification from the blood meals of common vampire bats (*Desmodus rotundus*)"

**Table of Contents:**

| **Table S1:** Information on where each common vampire bat blood meal sample was collected, and corresponding prey taxa analysed using metabarcoding | Page 2 |
| --- | --- |
| **Table S2:** Seeds used for *de novo* mtDNA contig assembly of prey taxa from common vampire bat bloodmeals | Page 2 |
| **Table S3:** Seeds used for *de novo* mtDNA contig assembly for common vampire bat blood meal samples with identified prey from *BLAST*-alignment step | Page 2 |
| **Table S4:** Seeds used for *de novo* mtDNA contig assembly for common vampire bat blood meal samples with no identified prey from *BLAST*-alignment step. Green: contig(s) assembled | Page 3 |
| **Table S5:** Seeds used for testing accuracy of *de novo* mtDNA contig assembly by using seeds from unidentified prey species. Green: contig(s) assembled | Page 4 |
| **Table S6:** Taxonomic identification of prey from common vampire bat bloodmeals based on mapping reads to COI barcode. The number of reads mapped to the COI barcode is shown for each bloodmeal sample | Page 5 |

**Table S1:** Information on where each common vampire bat blood meal sample was collected, and corresponding prey taxa analysed using metabarcoding

| **Sample** | **Site** | **Ecoregions** | **Year collected** | **Latitude** | **Longitude** | **Prey taxa from metabarcoding^1^** |
| --- | --- | --- | --- | --- | --- | --- |
| 25 | MDD130 | Amazon | 2012 | -1.300.944.999 | -7.065.340.994 | *Tapirus terrestris* |
| 29 | LMA4 | Coastal | 2010 | -1.266.557.638 | -7.666.797.959 | *Sus scrofa* |
| 54 | MDD130 | Amazon | 2009 | -1.300.944.999 | -7.065.340.994 | *Tapirus terrestris* |
| 70 | LMA6 | Coastal | 2012 | -11.06 | -77.46 | *Equus caballus* |
| 90 | LMA6 | Coastal | 2012 | -11.06 | -77.46 | *Equus asinus* |
| 94 | LMA10 | Coastal | 2012 | -1.158.834.583 | -7.727.840.376 | *Bos taurus* |
| 116 | MDD130 | Amazon | 2012 | -1.300.944.999 | -7.065.340.994 | *Tapirus terrestris* |
| 121 | MDD130 | Amazon | 2013 | -1.300.944.999 | -7.065.340.994 | *Gallus gallus and Tapirus terrestris* |

^1:^ Data retrieved from (Bohmann et al., 2018)

**Table S2:** Seeds used for *de novo* mtDNA contig assembly of prey taxa from common vampire bat blood meals

| **Species** | **Seed type** | **Database** | **Database number** | **Sequence length (bp)** |
| --- | --- | --- | --- | --- |
| *Bos taurus* | COI | BOLD | ACLB032-06 | 639 |
|  | mtDNA | GenBank | NC_006853.1 | 16338 |
| *Equus asinus* | COI | BOLD | GBMA100000-15 | 624 |
|  | mtDNA | GenBank | NC_001788.1 | 16670 |
| *Equus caballus* | COI | BOLD | GBGC18042-19 | 1542 |
|  | mtDNA | GenBank | NC_001640.1 | 16660 |
| *Gallus gallus* | COI | BOLD | ACLB009-06 | 639 |
|  | mtDNA | GenBank | NC_040970.1 | 16785 |
| *Sus scrofa* | COI | BOLD | GBMA4901-13_C01 | 653 |
|  | mtDNA | GenBank | NC_000845.1 | 16613 |
| *Tapirus terrestris* | COI | BOLD | ABRMM038-06-C01 | 657 |
|  | mtDNA | GenBank | AJ428947.1 | 16776 |

**Table S3:** Seeds used for *de novo* mtDNA contig assembly for common vampire bat blood meal samples with identified prey from *BLAST*-alignment step

| **Sample** | **Species** | **Seed type** | **Kmer size** | **Contigs** |
| --- | --- | --- | --- | --- |
| 54 | *Tapirus terrestris* | COI | 39 | Yes |
| 70 | *Equus caballus* | COI | 39 | Yes |
| 90 | *Equus asinus* | COI | 39 | No |
|  |  | COI | 27 | No |
|  |  | mtDNA | 39 | Yes |
| 94 | *Bos taurus* | COI | 39 | Yes |
| 121 | *Gallus gallus* | COI | 39 | Yes |

**Table S4:** Seeds used for *de novo* mtDNA contig assembly for common vampire bat blood meal samples with no identified prey from *BLAST*-alignment step. Green: contig(s) assembled

| **Sample** | **Species** | **Seed type** | **Kmer size** | **Contig output** |
| --- | --- | --- | --- | --- |
| 25 | *Bos taurus* | COI | 39 | No |
|  |  |  | 27 | No |
|  | *Equus asinus* | COI | 39 | No |
|  |  |  | 27 | No |
|  | *Equus caballus* | COI | 39 | No |
|  |  |  | 27 | No |
|  | *Gallus gallus* | COI | 39 | No |
|  |  |  | 27 | No |
|  | *Sus scrofa* | COI | 39 | No |
|  |  |  | 27 | No |
|  | *Tapirus terrestris* | COI | 39 | No |
|  |  |  | 27 | Yes |
| 29 | *Bos taurus* | COI | 39 | No |
|  |  |  | 27 | No |
|  |  | mtDNA | 39 | No |
|  |  |  | 27 | No |
|  | *Equus asinus* | COI | 39 | No |
|  |  |  | 27 | No |
|  |  | mtDNA | 39 | No |
|  |  |  | 27 | No |
|  | *Equus caballus* | COI | 39 | No |
|  |  |  | 27 | No |
|  |  | mtDNA | 39 | No |
|  |  |  | 27 | No |
|  | *Gallus gallus* | COI | 39 | No |
|  |  |  | 27 | No |
|  |  | mtDNA | 39 | No |
|  |  |  | 27 | No |
|  | *Sus scrofa* | COI | 39 | No |
|  |  |  | 27 | No |
|  |  | mtDNA | 39 | No |
|  |  |  | 27 | Yes |
|  | *Tapirus terrestris* | COI | 39 | No |
|  |  |  | 27 | No |
| 116 | *Bos taurus* | COI | 39 | No |
|  |  |  | 27 | No |
|  |  | mtDNA | 39 | No |
|  |  |  | 27 | No |
|  | *Equus asinus* | COI | 39 | No |
|  |  |  | 27 | No |
|  |  | mtDNA | 39 | No |
|  |  |  | 27 | No |
|  | *Equus caballus* | COI | 39 | No |
|  |  |  | 27 | No |
|  |  | mtDNA | 39 | No |
|  |  |  | 27 | No |
|  | *Gallus gallus* | COI | 39 | No |
|  |  |  | 27 | No |
|  |  | mtDNA | 39 | No |
|  |  |  | 27 | No |
|  | *Sus scrofa* | COI | 39 | No |
|  |  |  | 27 | No |
|  |  | mtDNA | 39 | No |
|  |  |  | 27 | No |
|  | *Tapirus terrestris* | COI | 39 | No |
|  |  |  | 27 | No |
|  |  | mtDNA | 39 | No |
|  |  |  | 27 | No |

**Table S5:** Seeds used for testing accuracy of *de novo* mtDNA contig assembly by using seeds from unidentified prey species. Green: contig(s) assembled

| **Sample** | **Prey identified** | **Seed species** | **Seed type** | **Contig identity** |
| --- | --- | --- | --- | --- |
| 25 | *Tapirus terrestris* | *Bos taurus* | COI |  |
|  |  | *Equus asinus* | COI |  |
|  |  | *Equus caballus* | COI |  |
|  |  | *Gallus gallus* | COI |  |
|  |  | *Sus scrofa* | COI |  |
| 29 | *Sus scrofa* | *Tapirus terrestris* | mtDNA |  |
|  |  |  | COI |  |
|  |  | *Bos taurus* | COI |  |
|  |  | *Equus asinus* | COI |  |
|  |  | *Equus caballus* | COI |  |
|  |  | *Gallus gallus* | COI |  |
|  |  | *Sus scrofa* | COI |  |
| 54 | *Tapirus terrestris* | *Bos taurus* | COI |  |
|  |  | *Equus asinus* | COI | *Tapirus terrestris* |
|  |  | *Equus caballus* | COI |  |
|  |  | *Gallus gallus* | COI |  |
|  |  | *Sus scrofa* | COI |  |
| 70 | *Equus caballus* | *Tapirus terrestris* | mtDNA | *Equus caballus* |
|  |  |  | COI |  |
|  |  | *Bos taurus* | COI |  |
|  |  | *Equus asinus* | COI | *Equus caballus* |
|  |  | *Gallus gallus* | COI |  |
|  |  | *Sus scrofa* | COI |  |
| 90 | *Equus asinus* | *Tapirus terrestris* | mtDNA |  |
|  |  |  | COI |  |
|  |  | *Bos taurus* | COI |  |
|  |  | *Equus caballus* | COI |  |
|  |  | *Gallus gallus* | COI |  |
|  |  | *Sus scrofa* | COI |  |
| 94 | *Bos taurus* | *Tapirus terrestris* | mtDNA | *Bos taurus* |
|  |  |  | COI |  |
|  |  | *Equus asinus* | COI |  |
|  |  | *Equus caballus* | COI | *Bos taurus* |
|  |  | *Gallus gallus* | COI | *Bos taurus* |
|  |  | *Sus scrofa* | COI | *Bos taurus* |
| 116 | NA | *Tapirus terrestris* | mtDNA |  |
|  |  |  | COI |  |
|  |  | *Bos taurus* | COI |  |
|  |  | *Equus asinus* | COI |  |
|  |  | *Equus caballus* | COI |  |
|  |  | *Gallus gallus* | COI |  |
|  |  | *Sus scrofa* | COI |  |
| 121 | *Gallus gallus* | *Tapirus terrestris* | mtDNA |  |
|  |  |  | COI |  |
|  |  | *Bos taurus* | COI |  |
|  |  | *Equus asinus* | COI |  |
|  |  | *Equus caballus* | COI |  |
|  |  | *Sus scrofa* | COI |  |

**Table S6:** Taxonomic identification of prey from common vampire bat blood meals based on mapping reads to COI barcode. The number of reads mapped to the COI barcode is shown for each blood meal sample

| **Sample** | **Class** | **Order** | **Family** | **Genus** | **Species** | **COI** |
| --- | --- | --- | --- | --- | --- | --- |
| 25 | NA | NA | NA | NA | NA | 0 |
| 29 | NA | NA | NA | NA | NA | 0 |
| 54 | Mammalia | Perissodactyla | Tapiridae | *Tapirus* | *Tapirus terrestris* | 2 |
| 70 | Mammalia | Perissodactyla | Equidae | *Equus* |  | 7 |
|  | Mammalia | Perissodactyla | Equidae | *Equus* | *Equus caballus* | 4 |
|  | Mammalia | Perissodactyla | Equidae | *Equus* |  | 1 |
| 90 | Mammalia | Perissodactyla | Equidae | *Equus* | *Equus asinus* | 1 |
| 94 | Mammalia | Artiodactyla | Bovidae |  |  | 2 |
|  | Mammalia | Artiodactyla | Bovidae | *Bison* | *Bison bonasus* | 1 |
|  | Mammalia | Artiodactyla | Bovidae | *Bos* |  | 13 |
|  | Mammalia | Artiodactyla | Bovidae | *Bos* | *Bos grunniens* | 1 |
|  | Mammalia | Artiodactyla | Bovidae | *Bos* | *Bos indicus* | 2 |
|  | Mammalia | Artiodactyla | Bovidae | *Bos* | *Bos primigenius* | 1 |
|  | Mammalia | Artiodactyla | Bovidae | *Bos* | *Bos taurus* | 15 |
|  | Mammalia | Artiodactyla |  |  |  | 1 |
| 116 | NA | NA | NA | NA | NA | 0 |
| 121 | Aves | Galliformes | Phasianidae | *Gallus* |  | 1 |
|  | Aves | Galliformes | Phasianidae | *Gallus* | *Gallus gallus* | 7 |
